# Siamese cat tyrosinase has enhanced proteasome degradation and increased cellular aggregation

**DOI:** 10.1101/2020.06.03.132613

**Authors:** Ingrid R. Niesman

## Abstract

Siamese cats are a notable example of a temperature-sensitive partial albinism phenotype. The signature color-pointing pattern is the result of an amino acid substitution – G302R – in the cysteine-rich domain of feline tyrosinase. The precise mechanism for the loss of tyrosinase enzyme activity due to this mutation is unknown.

**Objective:** We have used a cellular biology approach to begin unravel relationships between feline coloration, behavior and increased risk for feline cognitive dysfunction syndrome. GFP-fusion constructs of wild type domestic short hair tyrosinase and Siamese (G302R) tyrosinase generated to study cellular trafficking, degradation and the propensity for cellular aggregation.

**Data Description:** C-terminal GFP G302R expression has reduced Golgi localization, increased cytosolic fractions with reduced calnexin co-localization. N-terminal GFP constructs were retained in the ER, with little to no Golgi associated forms. C-terminal and N-terminal GFP G302R TYR is observed to have increased high molecular weight aggregation following proteasome inhibition.

## Introduction

Our objective is to understand if expression of mutant temperature-sensitive Siamese tyrosinase (TYR) is cytotoxic to cells. As TYR is expressed in neurons as well as melanocytes, toxicity over time can lead to unresolvable neuronal damage. We use classic cellular trafficking and proteasome inhibition to look for evidence of ER retention and up-regulation of the highly conserved unfolded protein response (UPR). The Siamese cat color-pointing pattern is one of the most recognizable patterns in the animal kingdom. First described in 2005 by Lyons, et al., [1] the distinct phenotype of Siamese is the result of a single point mutation (G>C) in the secreted endoplasmic reticulum (ER) glycoprotein, tyrosinase (TYR). In cooler areas TYR G302R is fully active, resulting in melanin production. In areas of normal temperature, such as in the brain, TYR G302R, has limited activity. Siamese mutant TYR has well-documented pleiotropic effects [2]. Early research reported differential effects on eye phenotypes [3] and in development of specific brain pathways [4–9] between random bred cats and Siamese. Humans also have comparable TYR point mutations that manifest a temperature-sensitive phenotype [10–12]. The mutant TYR R422Q misfolds at normal body temperature, after which it is retained in the ER where it can be toxic [13]. We understand little of the molecular basis for the temperature-sensitive phenotype of TYR in felines. If ER retentions and subsequent maladaptive UPR induction occur with the Siamese cat mutation, as in the human TYR R422Q mutation, then these cats represent a naturally occurring model to study feline cognitive dysfunction syndrome (fCDS).

## Materials and Methods

### Reagent list

The following antibodies were used for immunofluorescence and immunoblotting, (Abcam; CHOP mAb #2895T, GFP rabbit mAb #2956T Novus Biologicals; TGN38 mAb #NB300-575SS; Santa Cruz Biotechnology GAPDH mAb #sc-365062). Hela and Cos 7 cells were grown with DMEM-F12 (Gibco #11330-032). Pierce Coomassie Bradford reagent; #23200, LDS 4x sample buffer; #NP0007, 4-12% gels Criterion XT Bis-Tris; Biorad # 3450124, ALLN Sigma A6185; MG-132; Sigma 474787, Mirus Bio TransiT-LT1; #MIR2304, Agilent; QuikChange II #200523, Roche cOmplete protease inhibitor; #04693159001.

### Preparation of synthetic tyrosinase

Bioinformatics was used to extract relevant information about the feline genome. Full-length feline TYR (GeneID: 751100) is fully annotated (NC_018732.3), with five exons and four introns. mRNA (XM_0039992642.4) is 2418 base pairs and full length TYR protein (XP_0039992691.2) is 530 amino acids. Genscript synthesized the G320R TYR full-length cDNA sequence with EcoR1 and BAMH1 restriction sites engineered in for seamless cloning. The DSH wild type allele was derived by site directed mutagenesis). Full sequencing confirmed the single point change resulting in DSH clones. Digested TYR DNA was cloned into C2-EGFP (N-terminal EGFP-TYR) and N1-EGFP (C-terminal TYR-EGFP) with the stop codon removed by site directed mutagenesis [generous gifts of Dr. Christopher Glembotski]. All clones were sequence confirmed and large batch plasmid preparations isolated using spin column technology.

### Cell Culturing and Transfection Conditions

12 well tissue culture plates were seeded with 2 × 10^5^ cells per well 24 hours prior to transfection. For immunofluorescence, ethanol sterilized glass coverslips were placed in each well. 1mg of total cDNA was complexed with Mirus Bio TransiT-LT1 according to manufacturer’s instructions and dripped into individual wells. After 24 hours, GFP expression was evaluated by fluorescence microscopy to determine transfection efficiencies before continuing with experiments. Average transfection efficiency was 30-35% (data not shown) for all constructs. Empty GFP vectors served as controls. All experiments were performed at the 24-hour expression timepoint. For immunofluorescence experiments, coverslips were washed twice DPBS and immediately fixed with ice-cold methanol for 15 min at −20°C. They were washed in PBS with 0.1% Triton-X100 (washing buffer) and stored at 4°C. Coverslips were blocked with 5% FBS in washing buffer for 30 minutes at room temperature, followed by overnight incubations with primary antibodies at 4°C. Cells were washed and re-incubated for 2 hours rotating gently with appropriate secondary antibodies, washed and mounted with Vectashield-DAPI antifade reagent. Slides were imaged on an Olympus epifluorescence microscope or a Zeiss 710 confocal microscope.

### Western Blot Analysis

Cells were washed twice with DPBS and lysed with 75 μl lysis buffer (50mM Tris pH 7.5, 150mN NaCl_2_, 1% Triton-X100 and 01%SDS) with protease inhibitors added. Following sonication, lysates were centrifuged and supernatants normalized by Bradford protein assays. 20-30 μg of total protein were separated by electrophoresis on 4-12% gradient gels. Gels were transferred to PVDF membranes and blocked with 5% nonfat dry milk in TBS-Tween 20. Membranes were incubated overnight with primary antibodies at 4°C, washed and incubated with secondary antibodies at 1:5000 for two hours. Blots were imaged on an ImageQuant LAS 4000.

## Results

N-terminal constructs differ in localization to C-terminal constructs [14]. GFP tagged proteins at N-terminus and C-terminus were used in these studies to prevent biased analysis due to known issues arising from GFP folding [15]. By WB, C-terminal tagged fusion TYR shows a typical TYR pattern of higher molecular weight Golgi protein, mature glycosylated ER forms and a lower immature ER form. N-terminally tagged TYRs are not trafficked to the Golgi. Fluorescent analysis reveals a similar sub-cellular distribution. C-terminal constructs demonstrate clear ER-like localization with a large pool of GFP fluorescence perinuclear. A moderate pool of cytosolic GFP is evident, but the bulk of GFP aligns with membrane-bound organelles. N-terminal constructs are restricted to an ER-like domain, with limited cytosolic pools [16].

Subcellular co-localizations are altered in G302R TYR constructs [17]. Using Golgi disruption we see that DSH TYR is localized to stacks and G302R is not. The stacks become dispersed punctate large vesicles spread through the cytosol. C-terminal DSH TYR remains colocalized with the dispersed stacks. C-terminal G302R is no longer localized within the Golgi stacks. G302R TYR has reduced calnexin (CNX) colocalization in the ER, indicating G302R TYR is more rapidly translated, resulting in mixed populations of occupied N-linked glycosylation sites [18]. No construct aligns with Lamp1 as expected for non-pigmented cell types (25). Degradation is expected through proteasome mechanisms.

G302R TYR has increased ER retention and increased aggregation when proteasome degradation is inhibited [19]. We investigated the contributions of endoplasmic-reticulum-associated-protein-degradation (ERAD) pathways using a pan-calpain inhibitor (2.5mg/ml ALLN) and 26S proteasome inhibitor (5mm MG132). C-terminal G302R and N-terminal G302R displayed increased amounts of >150kD aggregates by WB when cells are treated for 6 hours. C-terminal G302R expression is retained in the ER where it causes ER dilation. Medium to large vesicles leak from the ER into the cytoplasm. We posit these vesicles are large cellular aggregates of G302R TYR that would normally be degraded due to misfolding events. In contrast, the bulk of DSH TYR is retained in the ER with limited cytosolic vesicles. A similar trend occurs when the C-terminal constructs are treated with 5mm MG132. The N-terminal constructs show a slight different pattern. When treated with ALLN, N-terminal DSH TYR is again retained in the ER where it colocalizes with CNX. G302R TYR dilates the ER and shunts much larger and more numerous GFP positive vesicles into the cytosolic domain. It is clear that G302R TYR is retained in the ER and is more prone to increased aggregation within the ER.

G302R TYR has enhanced degradation and ER stress [20]. Cycloheximide treatment of C-terminal constructs demonstrates that G302R is degraded at enhanced rates. Quality control mechanisms are highly sensitive to minimal changes in protein folding [13], leading us to conclude that G302R TYR frequently misfolds in the ER. When transfected cells are treated with 10mg/ml tunicamycin to elicit ER stress, G302R expressing cells show increased expression of the canonical ER stress marker; CHOP, again suggesting the G320R is retained in the ER and could prove cytotoxic.

**Figure 1.**
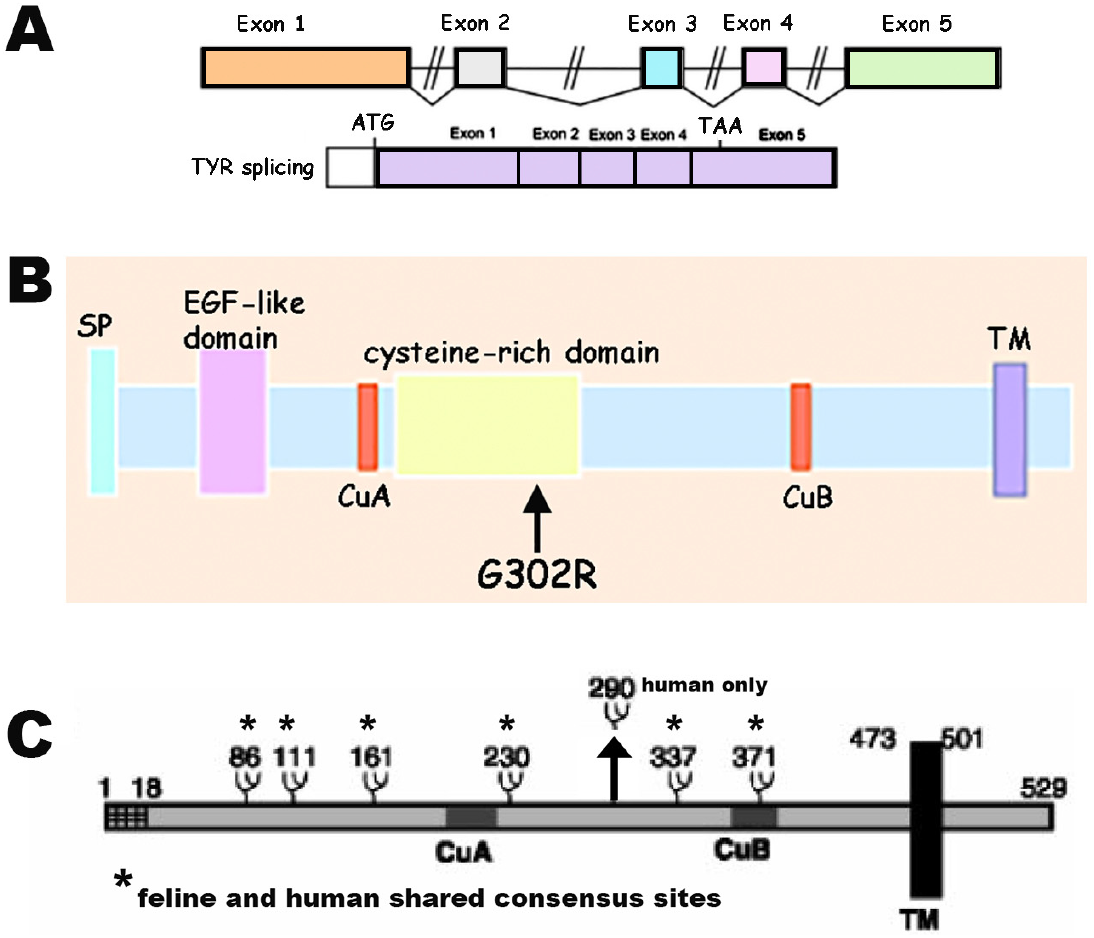
Bioinformatics of feline tyrosinase. **A**. Gene structure **B.** protein structure **C.** comparison of human and feline N-linked glycosylation sites

**Figure 2.**
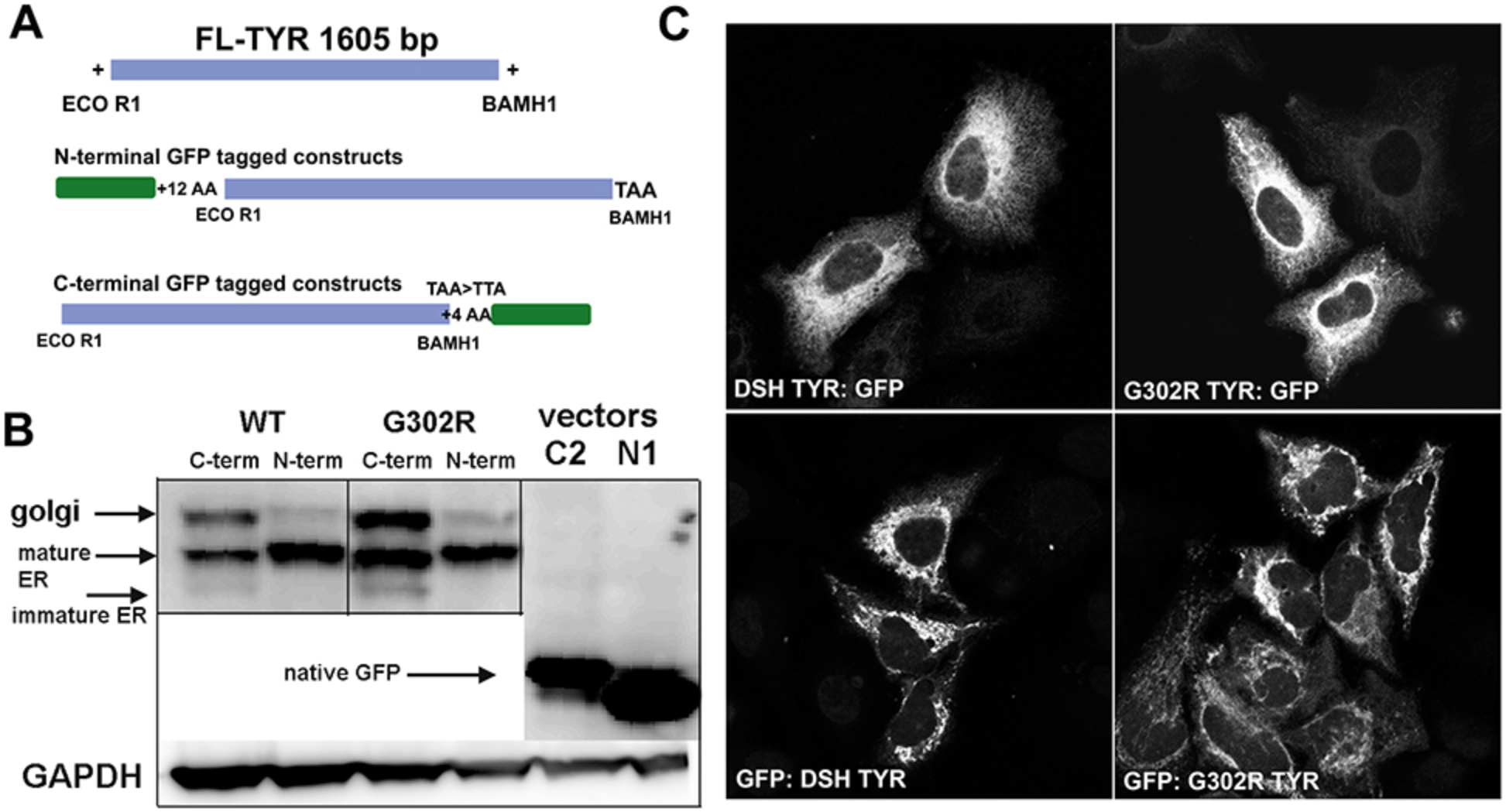
Expression of GFP fusion constructs. **A**. Construct designs **B**. WB analysis of WT-DSH and mutant G302R constructs in Hela cells after 24 hours **C**. Confocal microscopy analysis (Zeiss 710) of GFP fluorescence.

**Figure 3.**
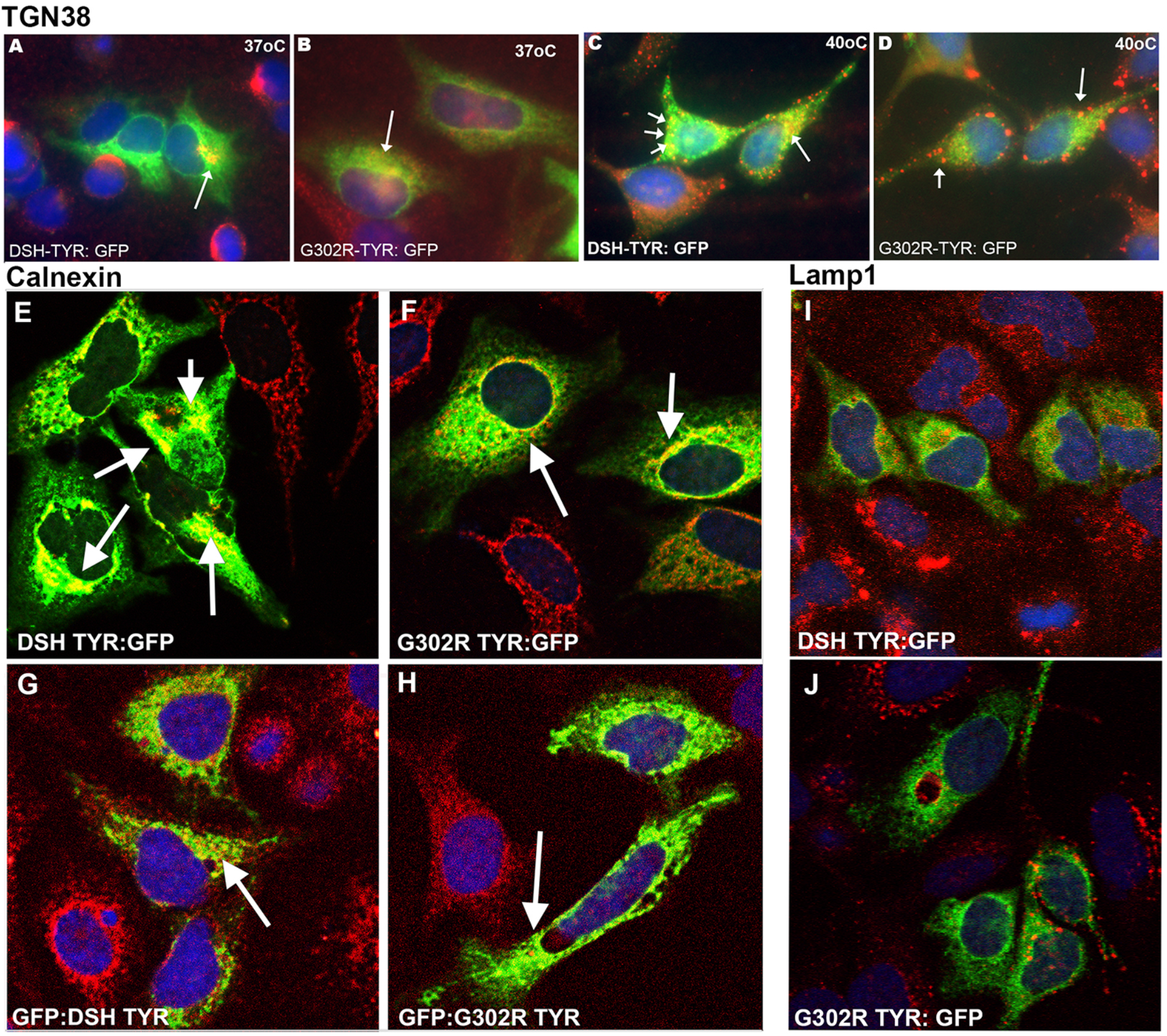
Colocalization with canonical cellular compartment markers. **A – D**. C-terminal constructs expresses in Hela cells stained with TGN38 (golgi) **E – H**. C-terminal and N-terminal constructs labeled with CNX **I – J**. C-terminal constructs labeled with lysosomal marker Lamp1.

**Figure 4.**
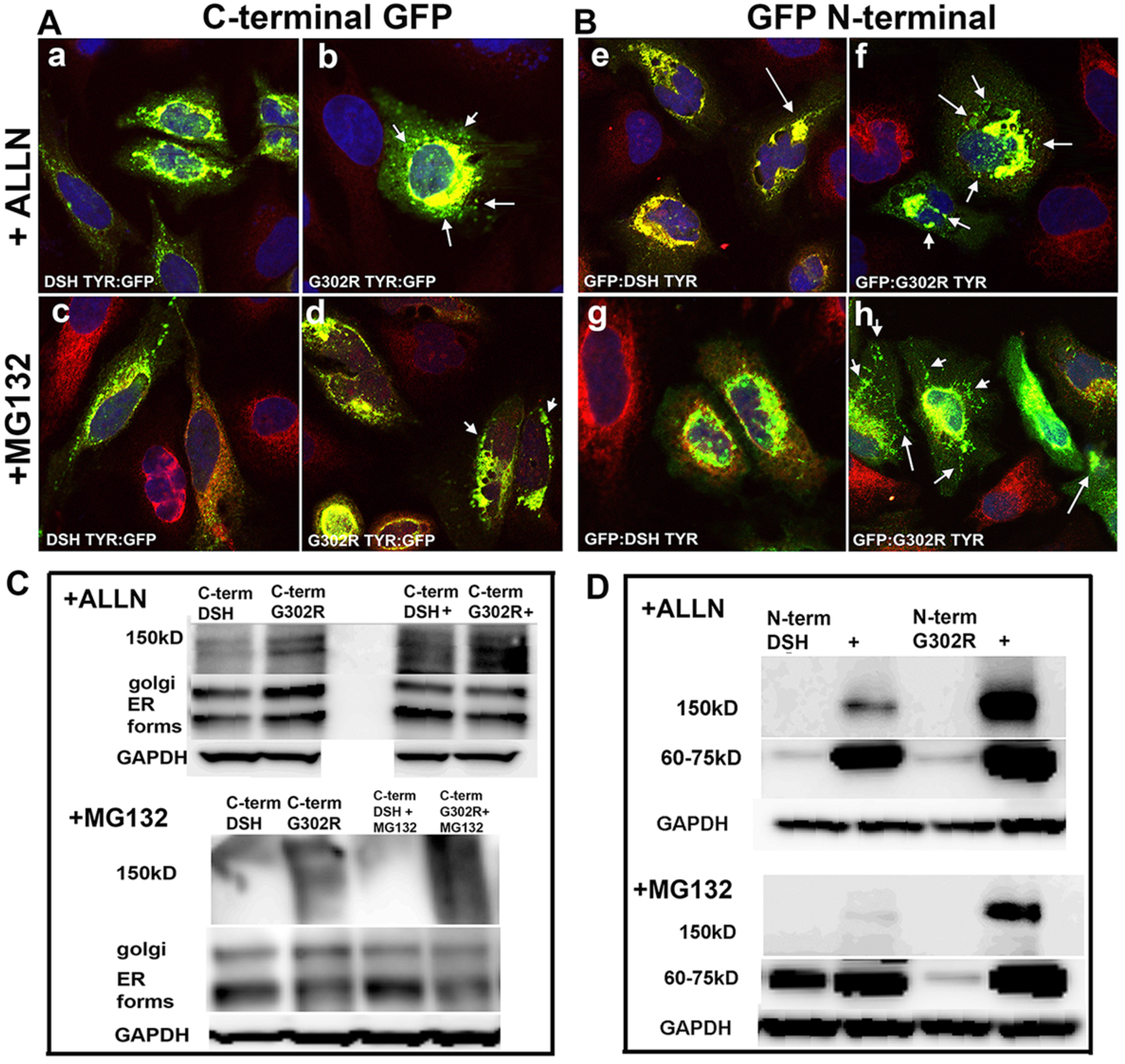
Mutant G302R constructs have enhanced aggregation when degradation is inhibited. **A**. C-terminal constructs stained with CNX **B.** N-terminal constructs stained with CNX **C.** C-terminal constructs WB following inhibition with ALLN and MG132 **D**. N-terminal constructs WB following inhibition with ALLN and MG132,

**Figure 5.**
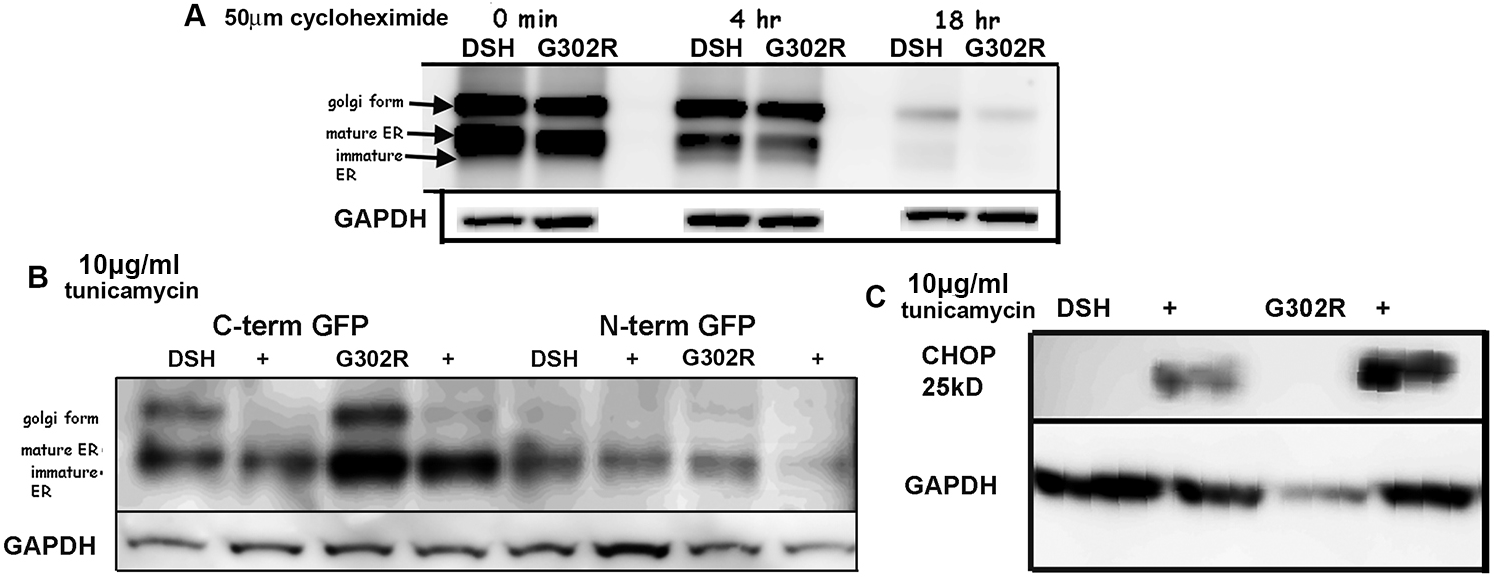
G302R TYR is degraded faster and has enhanced ER stress. **A**. CHX treatment timecourse of C-terminal constructs **B**. 6 hour Tunicamycin treatment WB **C**. WB of Tunicamycin treated cells and labeled with marker of UPR, indicating ER stress (CHOP).

## Discussion

These data are preliminary studies to determine if our construct designs expressed as expected. We carried out experiments in Hela or Cos7 cells first to compare our results to previously published models, knowing any effects of G302R would be highly leaky at a near permissive temperature (37°C). Cats have a normal body temperature of 40°C, considerably higher than normal tissue culturing temperatures. Cos7 cells provided increased expression but neither Hela or Cos7 cells held up for 24 hours at >39°C. Seeing even the modest effects of G302R over DSH is encouraging. We plan to move our studies to (1) feline kidney cells grown at 40°C and (2) isogenic feline iPSC lines of G302R and DSH TYR to study the aggregation and cytotoxicity in differentiated neurons directly at the non-permissive temperature. We are providing these results to encourage other researchers to pursue Siamese cats as a potential model of neurodegeneration.

## Abbreviations

Tyrosinase; TYR, calnexin; CNX, cycloheximide; CHX, N-Acetyl-Leu-Leu-Norleu-al, N-Acetyl-L-leucyl-L-leucyl-L-norleucinal; ALLN, Z-L-Leu-D-Leu-L-Leu-al; MG-132

## Declarations

### Ethics approval and consent to participate

All regulatory policies regarding use of human cell lines and recombinant DNA are approved by SDSU IBC, protocol # 19-01-002N.

### Consent for publication

Not Applicable

### Availability of data and materials

The data described in this manuscript can be freely and openly accessed on ***[FigShare]***. Please see **Table 1** and references for details and links to the data.

**Table 1.** Overview of data files/data sets.

| Label | Name of data file/data set | File types<br>(file extension) | Data repository and<br>identifier (DOI or accession<br>number) |
| --- | --- | --- | --- |
| Figure 1 | Bioinformatics of feline tyrosinase | .jpg | https://doi.org/10.6084/m9.figshare.12331220.v1 |
| Figure 2 | Expression of GFP fusion constructs | .jpg | https://doi.org/10.6084/m9.figshare.12331223.v1 |
| Figure 3 | Colocalization with canonical cellular compartments | .jpg | https://doi.org/10.6084/m9.figshare.12331226.v1 |
| Figure 4 | Mutant G302R constructs have enhanced aggregation with degradation is inhibited | .jpg | https://doi.org/10.6084/m9.figshare.12341030.v1 |
| Figure 5 | G302R tyrosinase is degraded faster and has enhanced ER stress | .jpg | https://doi.org/10.6084/m9.figshare.12331232.v1 |

### Competing interests

No completing interests

### Funding

All work on this project was self-funded by Ingrid Niesman in a donation to the SDSU Research Foundation.

### Authors’ contributions

**I.R.N. developed the project, designed the constructs, directed the experimental work, and wrote the manuscript.**

## Acknowledgements

This work would not have been possible without the generous advice and scientific input from Dr. Christopher Glembotski – Distinguished Professor of Biology at SDSU. He allowed me to use his lab and equipment to begin this work.

## Notes

https://doi.org/10.6084/m9.figshare.12331223.v1

https://doi.org/10.6084/m9.figshare.12331226.v1

https://doi.org/10.6084/m9.figshare.12341030.v1

https://doi.org/10.6084/m9.figshare.12331232.v1

https://doi.org/10.6084/m9.figshare.12331220.v1

### Competing Interest Statement

The authors have declared no competing interest.

